# Preliminary Evaluation of the Utility of Deep Generative Histopathology Image Translation at a Mid-Sized NCI Cancer Center

**DOI:** 10.1101/2020.01.07.897801

**Authors:** Joshua J. Levy, Christopher R. Jackson, Aravindhan Sriharan, Brock C. Christensen, Louis J. Vaickus

## Abstract

Evaluation of a tissue biopsy is often required for the diagnosis and prognostic staging of a disease. Recent efforts have sought to accurately quantitate the distribution of tissue features and morphology in digitized images of histological tissue sections, Whole Slide Images (WSI). Generative modeling techniques present a unique opportunity to produce training data that can both augment these models and translate histologic data across different intra-and-inter-institutional processing procedures, provide cost-effective ways to perform computational chemical stains (synthetic stains) on tissue, and facilitate the creation of diagnostic aid algorithms. A critical evaluation and understanding of these technologies is vital for their incorporation into a clinical workflow. We illustrate several potential use cases of these techniques for the calculation of nuclear to cytoplasm ratio, synthetic SOX10 immunohistochemistry (IHC, sIHC) staining to delineate cell lineage, and the conversion of hematoxylin and eosin (H&E) stain to trichome stain for the staging of liver fibrosis.

## 1 INTRODUCTION

The field of histopathology is the study of disease using morphological and spatial distributions of tissue features observed under a microscope as performed by a board-certified pathologist. Tissue is biopsied or excised, fixed via a combination of formalin and heat processing, embedded in paraffin to create formalin-fixed and paraffin-embedded tissue blocks (FFPE). The tissue is then sectioned via a microtome into thin sheets (typically 2μM-8μM in thickness), and chemically stained with various reagents (most commonly hematoxylin and eosin, H&E). Slides and tissue blocks are typically stored for at least 10 years (depending on legal and institutional regulations), although there is no effective limit to their shelf life beyond slow bleaching of dyes and degradation of DNA/RNA and many academic institutions retain all tissue indefinitely. Despite advancements in “gold-standard” scoring metrics for tissue examination, some diagnoses have a high inter-observer disagreement between pathologists.. Like other medical professionals, pathologists are increasingly overworked and reporting high levels of burnout (Miller & Brown, 2018) which can significantly impact turn-around-time (TAT) and in severe cases, diagnostic accuracy. Moreover, maintaining a CLIA-certified (Raab, 2000) histopathology laboratory capable of performing special stains and immunohistochemistry (IHC) is expensive and requires specialized technologists, licenses, instruments and reagents. Thus, there is a great need for the development of accurate, interpretable, quantitative medical decision aid algorithms and workflows (Levy et al., 2020). In this article, we evaluate the application of generative, quantitative techniques in a variety of common scenarios in the application of digital pathology to current analogue workflows.

Generative modelling techniques aim to produce partially or wholly synthetic data that has a similarly high fidelity as the original or target data source. In some cases, data produced by these generative models can be used to enhance the performance of a variety of downstream prediction or inference tasks without the need to acquire additional, expensive, expert annotated data. These modelling techniques are useful in a wide variety of domains, including inferring ties in social networks for modelling the spread of healthy behaviors, robust translation for hundreds of languages, and the imputation of genomics data for the inference of more favorable disease prognosis as it relates to therapy choices (Lotfollahi et al., 2019; O’Malley, 2013; Young et al., 2018).

An important application of these generative techniques is the use of deep learning in medical imaging. Deep learning approaches are data-driven computational heuristics that “learn” to specify a large set of nonlinear interactions between predictors to understand their relationship with clinical outcomes of interest through the use of artificial neural networks (ANN) (Krizhevsky et al., 2012). ANN are comprised of layers of nodes that represent levels of abstraction of the data, where nodes aggregate weighted information from the previous layer of nodes, transform the data, and pass it to downstream layers of nodes to characterize the tissue image. We focus on a subclass of these methods, generative adversarial networks (GANs).

GANs simultaneously train two ANN. The first network generates an image of, for instance, histological tissue, from a latent data distribution (generator), and the other ANN discriminates whether the supplied image is “real” or “fake” (discriminator/critic). The number of publications on GANs in medical imagery has rapidly increased in recent years, with topics spanning medical image segmentation, nucleus detection, style translation, and upsampling of images (Yi et al., 2019).

A growing collection of studies have used GANs to synthetically stain images of histological tissue sections, which can save institutions time and money (both in reagents and technologists’ time) (Bayramoglu et al., 2017; Borhani et al., 2019; De Biase, 2019; Lahiani et al., 2018; Quiros et al., 2019; Rana et al., 2018; Rivenson, Liu, et al., 2019; Rivenson, Wang, et al., 2019; Xu et al., 2019). GAN models have also been used to remove artificial and natural discolorations in images of stained histological tissue sections that could perturb deep learning analyses (Bentaieb & Hamarneh, 2018; Ghazvinian Zanjani et al., 2018; Pontalba et al., 2019). Other studies have sought to use GANs to generate synthetic training data to increase the generalizability of deep learning histopathology models (Wei et al., 2019). A few studies used deep generative techniques to derive nucleus masks without the use of physician supplied annotations (Bug et al., 2019; Gadermayr et al., 2019; Hollandi et al., 2019; Mahmood et al., 2018).

Traditional unconditional GANs generate data by using the discriminator to estimate and match the distribution of the generated data to the real data. Conditional GANs such as Pix2Pix (Isola et al., 2018) are conditioned on the original labelled data while attempting to directly match the target image. This requires the pairing of images to learn the mapping from source to target domain (Figure 1a). In histopathology, perfectly registered paired images of differentially stained tissue can often be difficult to acquire (even when stereotactic sections are only separated by 5μM) and therefore require accurate annotation and registration, in which the images must be warped, rotated and translated to most accurately match the source domain. These registrations are not entirely accurate and assume that the matched tissue has no artifacts present (Borovec et al., 2019). Additionally, the computational resources required to register high resolution WSI with up to 80,000×80,000 pixels per color channel can be exceedingly high. Another GAN architecture resolves these issues: unconditional image-to-image translation models such as CycleGAN (Zhu et al., 2018) and UNIT (Zhu et al., 2018). In addition to distributional matching of the real and synthetic target and source domains, CycleGAN models learn to recapitulate the source/target domain after using a generator to construct the target/source image and another generator to reconstruct the original image (Figure 1b), which obviates the need to include matched images. A mathematical description of these two deep learning techniques have been included in the appendix.

**Figure 1:**
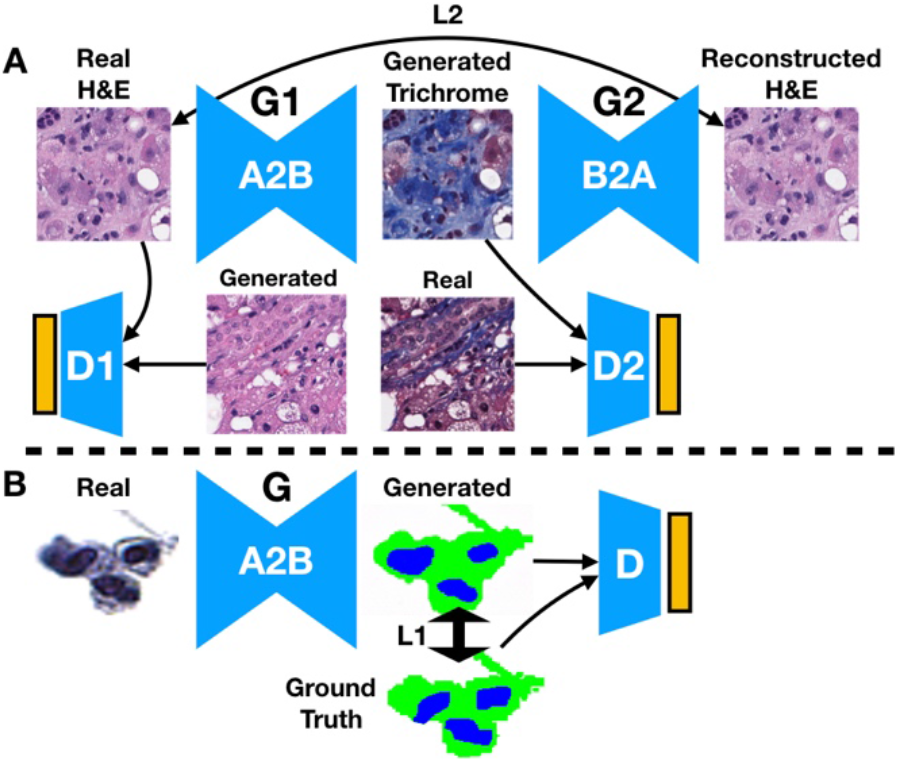
Illustration of (A) CycleGAN and (B) Pix2Pix Deep Learning Techniques. (A) Trichrome stain is generated from an H&E input image, which is in turn is used to generate a reconstruction of the original H&E image. The reconstructed image is compared to the original via a reconstruction loss: the distribution of generated stains are compared to the distribution of original stains via a discriminator. The same process is used when converting trichrome to H&E. (B) The urine segmentation mask is generated from the real image. Real and generated distributions are parameterized and compared both via the discriminator and directly between paired generated masks and ground truth segmentation masks.

In light of the successes of these GAN techniques in other domains, we sought to explore their applications to the WSI’s generated during clinical operations at the Dartmouth-Hitchcock Medical Center Department of Pathology and Laboratory Medicine (DHMC-PLM, a mid-sized National Cancer Institute Cancer Center, NCI-CC). In this study, we present preliminary results for the application of the CycleGAN and Pix2Pix techniques in calculating the nuclear to cytoplasm ratio of cells in urine cytology specimens, the conversion of H&E stains to predicted SOX10 immunohistochemistry, and the trichrome staining of liver tissue for fibrosis analysis.

## 2 METHODS

### 2.1 Data Preparation

Liver core needle biopsies obtained by various approaches were fixed in formalin and processed into FFPE tissue blocks; tissue sections were cut at 5μm thickness and stained with H&E and trichrome stains on adjacent levels, then scanned using the Leica Aperio-AT2 scanner at 20x magnification. The resulting images were stored natively as SVS files and then converted to NPY and ZARR files for preprocessing using PathFlowAI (Levy et al., 2020). An SQL database was created containing patches with 95% or greater non-white space. We extracted 500,000 and 290,000, unpaired, 256 pixel x 256 pixel subimages from 241 H&E and Trichrome stained WSI respectively as training data for a CycleGAN model. Seventy-five percent of the liver WSI (n=178 specimens with both H&E and trichrome WSI) were used to train the model, while 25% of the data was reserved for testing (n=63 specimens with both H&E and trichrome WSI).

For the urine cytology dataset, a total of 217 ThinPrep^®^ urine cytology slides were collected and scanned at 40x magnification using a Leica Aperio-AT2 scanner to 80K x 80K SVS files. These files were extracted to TIF images and resized to 40K x 40K. Via a process of background deletion and connected component analysis (Vaickus et al., 2019) we created libraries of cells from each of the WSI and randomly sampled 10,936 cell images from the 25×10^6^ extracted cellular objects. Manually annotated and algorithmically (AutoParis) derived segmentation masks delineating pixel-by-pixel areas of the nucleus, cytoplasm and background were paired with the original cell images (Vaickus et al., 2019). The cell images were separated into 60% training, 20% validation and 20% testing sets for Pix2Pix.

For the SOX10 IHC, an unpaired dataset containing a total of 15,000 H&E and 15,000 IHC, 256 pixel x 256 pixel subimages of skin and lymph node histology were acquired to train a CycleGAN model. Next, 30,000 paired H&E and IHC patches were registered to each other using an alignment technique that constructs a coarse pose transformation matrix to perform an initial alignment of the tissue, then dynamically warps and finetunes the IHC to the respective H&E tissue. Alignments with significantly poor warping were filtered out to yield 15,000 paired images for training a Pix2Pix model. The Pix2Pix IHC model data was split into 60% training, 20% validation and 20% testing. The CycleGAN model data was split into 60% training and 20% validation sets and shared the same test set with the Pix2Pix model. An additional 20 melanoma and 18 nevus cases (split into 224 x 224 pixel subimages) were processed for subjective grading of Pix2Pix and CycleGAN model outputs.

### 2.2 Analytic Approach

A CycleGAN model was fit on the liver dataset with the purpose of converting H&E stained images into trichrome stained images. We prioritized the ability of the model to identify pronounced fibrous tissue (portal tracts, large central veins, cirrhosis: highlighted in blue by the trichrome stain). Accordingly, highly blue patches were upsampled (as determined by blue color channel thresholds) and the reconstruction loss of trichrome stained tissue of the CycleGAN was upweighted to 0.8 versus H&E reconstructions (a process we are still fine-tuning). The model was trained for a total of 6 epochs and then the resulting model was utilized to construct entire synthetic trichrome WSI from H&E WSI in the test set. As a preliminary analysis, these synthetic images were qualitatively evaluated for their fidelity to the original trichrome (understain, equivalent, overstain) and then a pathologist scored the synthetic and real trichromes for the presence or absence of advanced fibrosis (bridging fibrosis or cirrhosis).

Pix2Pix was used to predict nucleus and cytoplasm segmentation masks for cells contained in urine cytology specimens. The Pix2Pix model was trained for 200 epochs using the default settings of a publicly available repository and then evaluated on a ground truth, hand-annotated test set. Pixel-level accuracy, sensitivity and specificity of each segmentation assignment (nucleus, cytoplasm, background) to the original ground truth was calculated and then the nucleus to cytoplasmic ratio (NC) was compared using the R^2^ correspondence. We utilized 1000-sample nonparametric bootstrap estimates to derive 95% confidence intervals for each of these estimates.

For the SOX10 dataset, both CycleGAN and Pix2Pix models were fit to the synthetic IHC dataset and each model was trained for approximately 150 epochs. These models were then utilized to convert H&E images into SOX10 IHC stained images. Color deconvolution algorithms were able to decompose both the real and generated IHC stained images into SOX10-positive (via 3,3 ‘-Diaminobenzidine (DAB) color deconvolution) and SOX10-negative (via hematoxylin color deconvolution) binary masks using color thresholding. Since there was imperfect registration between the H&E and IHC images, the resultant binary masks for each of the stains could not be directly compared. Instead, the area of the SOX10 positive and SOX10 negative stains were compared between the real and generated images across the test set via 1000-sample nonparametric bootstrap estimates of the 95% confidence intervals of the correlation coefficient between the area of real and generated positive and negatively stained tissue.

The architectures for the generators for both the CycleGAN and Pix2Pix models utilized residual neural network blocks (He et al., 2015).

## 3 PRELIMINARY RESULTS

Herein, we present our initial results from the application of the aforementioned tasks:

### 3.1 Cytology Segmentation

The NC ratio of urothelial cells is a metric that is estimated by pathologists and considered in combination with subjective measures of atypia to screen urine cytology specimens for urothelial carcinoma (bladder cancer) according to the gold standard Paris System for Urine Cytology (Barkan et al., 2016). The authors recently published a hybrid morphometric and deep learning approach to automating the Paris System utilizing a series of specialists semantic segmentation networks for NC ratio calculation requiring thousands of hand annotated images (Vaickus et al., 2019). Improving the quality and automation of cell compartment segmentation could provide significant performance gains to this and other automated techniques for the performance of cytological cancer screening tests (Layfield et al., 2017; Wang et al., 2019). Pix2Pix, utilizing a rather small training set, achieved remarkable segmentation performance, yielding an macro-accuracy of 0.95 and an R^2^ value of 0.74±0.019 between the ground truth and predicted NC ratios across the test set (Table 1; Figure 2).

**Figure 2:**
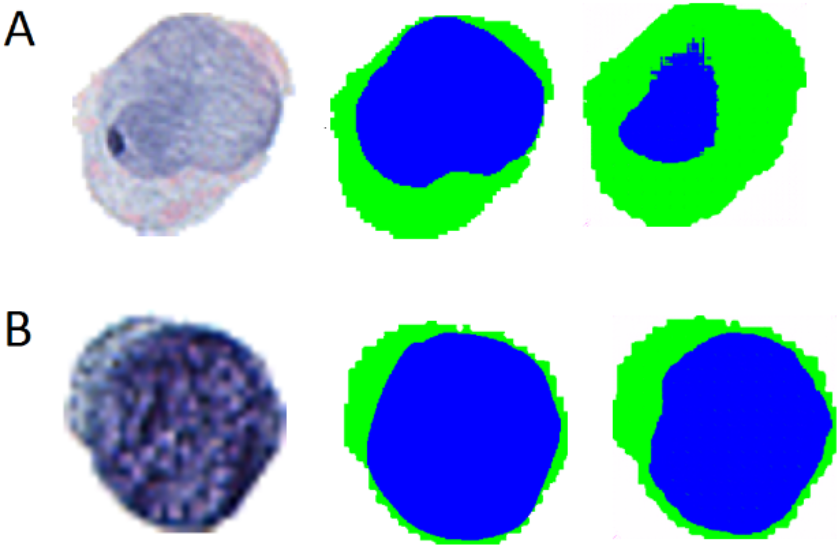
Examples of original cell images (left), ground truth segmentation masks (middle), and predicted segmentation masks (right) for: (A) poor segmentation and (B) excellent segmentation.

**Table 1:** Performance of Pix2Pix Urine Cell Nuclear and Cytoplasm Segmentation.

| Annotation Type | Accuracy ± SE | Sensitivity ± SE | Specificity ± SE |
| --- | --- | --- | --- |
| Background | $0.98 \pm 0.01$ | $0.98 \pm 0.01$ | $0.98 \pm 0.01$ |
| Cytoplasm | $0.93 \pm 0.05$ | $0.91 \pm 0.07$ | $0.95 \pm 0.05$ |
| Nucleus | $0.94 \pm 0.05$ | $0.85 \pm 0.16$ | $0.96 \pm 0.05$ |

### 3.2 Synthetic IHC

SOX10 is a nuclear transcription factor that is used in IHC for the identification of cells with melanocytic / neural crest origin (melanocytes, melanoma, etc). Immunohistochemical stains generate a distinctive brown color (DAB) on counterstained (hematoxylin) tissue sections allowing for relatively simple color deconvolution. Previous deep learning approaches have been able to utilize mappings between H&E and IHC tissue to learn antibody driven features in the H&E that may be correspondent to the separation of tumorous/non-tumorous tissue (Bulten & Litjens, 2018; Mohamed et al., 2013, p. 10; Willis et al., 2015). The ability of a deep learning model to predict the expression of DAB immunohistochemistry for a nuclear transcription factor (SOX10) from an HE stained image was first demonstrated by a resident pathologist in our research program (Christopher Jackson, MD) and presented at the 2019 meeting of the American Society of Dermatopathology (*full manuscript currently in review*) (Jackson, 2019).

Utilizing CycleGAN, we found that the area of SOX10 positive and negative staining was weakly associated between the predicted and true IHC stains (Table 2; Figures 3–4). However, when we trained on pairs of imperfectly registered images using Pix2Pix, we found much stronger correlations in the area of SOX10 staining between predicted and true IHC stains (Table 2; Figure 4). We also investigated each algorithm’s ability to identify melanocytic tissue for subjective analysis by pathologists and residents, in which the superior performance of the Pix2Pix model was consistently noted (Table 2; Figure 3–4).

**Figure 3:**
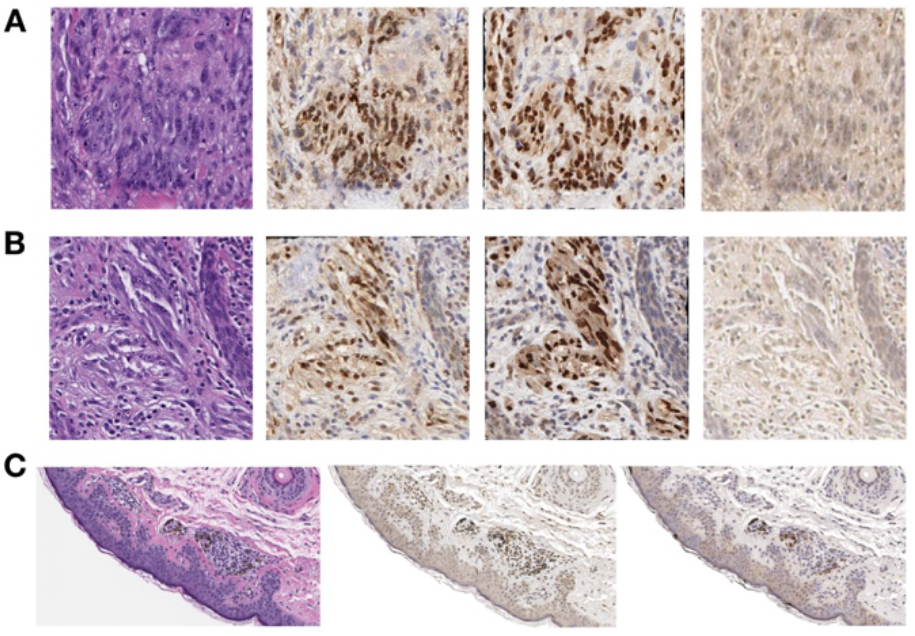
Examples of Synthetic IHC Staining Technique on: (A-B) Select image patches, organized by original H&E (left), Pix2Pix generated IHC (left-center), True Registered IHC (right-center), and CycleGAN generated IHC (right); (C) Pix2Pix (center) and CycleGAN (right) IHC images generated from large section of H&E stained tissue (left).

**Figure 4:**
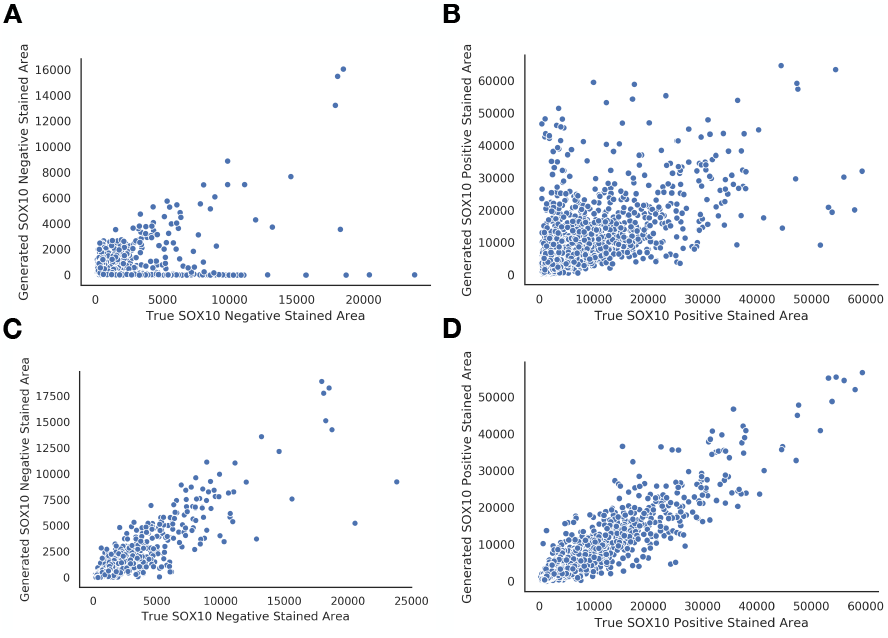
Breakdown of true versus predicted SOX10 stained area for: (A, B) CycleGAN; (C, D) Pix2Pix; (A, C) SOX10 Negative Staining; (B, D) SOX10 Positive Staining

**Table 2:** Comparison of CycleGAN versus Pix2Pix performance on synthetic SOX10 IHC staining.

| Method | Positive SOX10 Stained Area Predicted vs True PearsonR ± SE | Negative SOX10 Stained Area Predicted vs True PearsonR ± SE |
| --- | --- | --- |
| CycleGAN | 0.66±0.021 | 0.39±0.065 |
| Pix2Pix | 0.93±0.0061 | 0.90±0.013 |

### 3.3 Synthetic Trichrome Staining of Liver Tissue

Non-alcoholic steatohepatitis (NASH) is characterized by steatosis and chronic inflammation causing progressive liver injury and fibrosis in patients where no alcoholic, genetic, metabolic, or medication-based causes for hepatitis have been identified (Masugi et al., 2017). Progressive NASH can lead to cirrhosis with morbidities including ascites, sepsis, coagulopathies, and nutritional deficiencies, and a markedly increased risk of hepatocellular carcinoma (HCC). End stage cirrhosis requires transplantation which has a tremendous financial impact on the healthcare system and is associated with substantial patient morbidity and mortality. Fibrosis progression is typically assessed via liver biopsy. After core needle or wedge biopsies of the liver are obtained, pathologists score the tissue for features of NASH (percentage steatosis, presence of inflammatory cells and ballooning hepatocytes) using an H&E stain, then stage the degree of fibrosis with a trichrome stain, on which collagen (fibrous tissue) is highlighted blue. Here, we used CycleGAN to convert an H&E stain to a synthetic trichrome stain on small image patches and combined these patches to recapitulate the entire WSI (Figure 5). In visual assessment of gross trichrome stained area by a pathologist, the model showed a propensity for overcalling fibrosis with 58% deemed subjectively overstained, 37.5% deemed subjectively equivalent to the real trichrome, and 4.1% deemed subjectively understained. When a pathologist staged each synthetic and real trichrome stain for the presence or absence of advanced fibrosis (bridging fibrosis, cirrhosis), the accuracy of the synthetic trichromes was 79%, the sensitivity was 100% and the specificity was 67% (n=24, Table 3).

**Figure 5:**
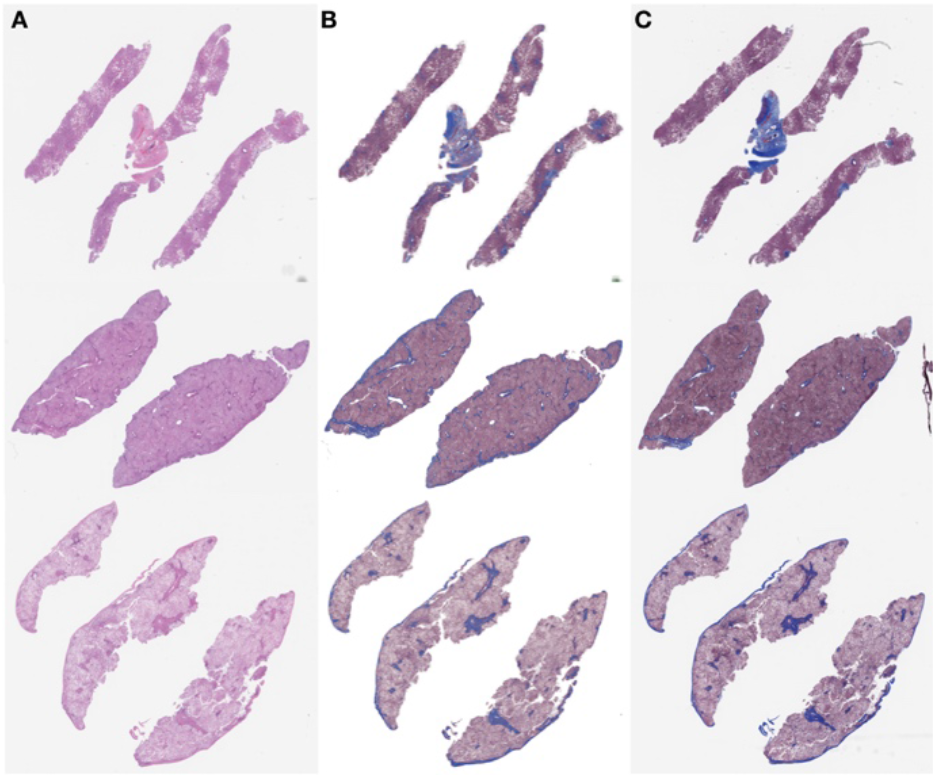
Examples of liver tissue specimen.(A) Original H&E stained tissue; (B) generated trichrome stain; (C) original matched trichrome stain

**Table 3:** Confusion matrix for synthetic trichrome in pathologist scoring of advanced fibrosis (bridging fibrosis, cirrhosis) versus non-advanced fibrosis.

|  |  |  |  |  |
| --- | --- | --- | --- | --- |
| N = 24 | Real + | Real - | Accuracy | 0.79 |
| Synth + | 11 | 6 | Sensitivity | 1.00 |
| Synth - | 0 | 12 | Specificity | 0.67 |

## 4 DISCUSSION

In this study, we illustrated a few potentially useful applications of CycleGANs and Pix2Pix in the clinical workflow of a pathology department at a midsized NCI Cancer Center. We found these synthetic staining and segmentation techniques to yield satisfactory performance in some situations, which we will continue to evaluate with larger datasets in the future. GANs are relatively difficult to train, partially in light of the non-convergence of the model’s objective function after training for a long period of time. The oscillation of the model loss during training reflects the dynamic min-max objective function between the generator and discriminator. For incorporation into the clinical workflow, these technologies will have to be packaged in ways that are accessible to the clinical researcher and clinician through simple GUIs and common workflow specifications (Amstutz et al., 2016) as well as provide for robust model validation.

We noted that Pix2Pix was able to accurately differentiate nuclei from cytoplasm in cells from urine cytology specimens. However, we noticed instances where the model both over and under-called nuclear area. This may be remedied by training the model on larger datasets with a more diverse sampling of nuclear and cytoplasmic morphologies.

Automated conversion of H&E to IHC presents an exciting opportunity to directly train models off of objective molecular targets which may decrease the bias present in physician annotations. Our preliminary results highlight the importance of using registration to assist with the accurate conversion of H&E to IHC. Regardless, we plan to investigate more robust measures of tissue similarity between ground truth and predicted stains (e.g. correspondence in circular objects detected using SURF and SIFT features) (Bay et al., 2006; Lowe, 2004). Current image registration techniques are also computationally intensive and imperfect (in this study it took 5 days to register 12 HE WSI to their respective IHC WSI). Further improvements in registration accuracy are likely to improve the synthetic IHC staining accuracy and we intend to invest significant research time in this area.

Our initial attempt at training a liver H&E to a synthetic trichrome staining model led to the gross under-prediction of liver fibrosis. We remedied this deficiency by supplying more images of highly fibrous tissue areas in the training set and by weighting to emphasise the reconstruction of fibrotic tissue. As a result of this, the overall accuracy of synthetic trichromes increased, but, perhaps predictably, led to a moderate degree of overestimation of fibrous tissue. Accordingly the sensitivity of the model for advanced fibrosis was high but the specificity was subpar. This result would likely be acceptable for a screening test, but given that the consequences of both over and under-calling liver fibrosis level are severe, synthetic trichromes for liver fibrosis staging would require very high sensitivity and specificity to be clinically usable. Further tuning of the model hyperparameters, architectures and/or the application of other generative techniques will be necessary to achieve better accuracy (Xu et al., 2019).

Given the superiority of the Pix2Pix in the synthetic IHC task, we intend to attempt to register the H&E and trichrome training images (however imperfectly). Considering that in every case the HE section is only stereotactically separated from the trichrome section by a 5 μM, this approach may prove successful, however, it is largely reliant on expert sectioning and placement of tissue slices by our histotechnologists. Regardless, our results provide a framework to improve upon these synthetic staining techniques for incorporation into the clinical workflow.

In light of these investigations, we find that generative models (CycleGAN, Pix2Pix) are well-positioned to supplement engineering solutions that are being developed to improve the efficiency and accuracy of histopathological diagnosis. Generally, clinicians at our institution are most interested in digital aids that either cut-down on cost and time to render a diagnosis or that automate tedious, error-prone tasks (such as IHC quantitation). These concerns are well met by our initial investigations. In the future, we will consider applications of generative models for the detection of cells for our clinical workflow, for standardizing stains from a collection of different institutions by translating to a common synthetic stain, and for superresolution techniques that may enable high-precision digital pathology from low fidelity source material.

## 4 CONCLUSIONS

In this study, we assessed the ability of deep learning image-to-image translation models to perform nucleus/cytoplasm segmentation in cells from urine cytology specimens, and synthetic trichrome and IHC staining on liver and skin WSI respectively. These initial investigations provide further evidence in favor of the incorporation of generative models into the core clinical workflow of a hospital pathology service. These methods may reduce the costs and time associated with chemical staining and manual image annotations while providing unbiased means for the molecular assessment of tissue. We will continue to assess additional applications of these technologies on new use cases with more robust measures of concordance and larger repositories of data.

## ACKNOWLEDGEMENTS

This work was supported by NIH grant R01CA216265 and a grant from the DHMC Norris Cotton Cancer Center, Cancer Fellows Program. JL is supported through the Burroughs Wellcome Fund Big Data in the Life Sciences at Dartmouth.

## APPENDIX

### 1 Mathematical Description of CycleGAN and Pix2Pix Objective Functions

GANs are tasked with learning a source/target domain *Y* from latent noise vector *Z* via mapping *G*. The loss function for a GAN is specified by:

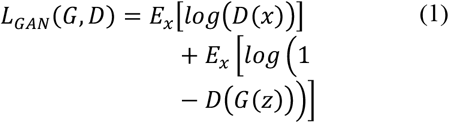

The ideal generator is acquired through alternating updates to the generator and discriminator parameters:

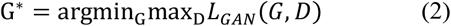

The objective is minimized with respect to the generator parameters to maximize the discriminator’s output for generated data D(G(z)) to attempt to fool the discriminator, which aims to maximize the separation between the real and generated data, as parameterized by the discriminator. The loss is maximized with respect to the discriminator’s parameters. GANs can be difficult to train, thus alterations to the objective such as the Wasserstein distance and gradient penalties have been introduced.

The goal of Image-to-Image translation is to learn a mapping from source domain/observed image *X* to a target domain *Y* via a generator *G*: *X* → *Y*.

The generator for both of the Pix2Pix and CycleGAN models compress input data *X* into a latent subspace *Z* by subsequent applications of convolutional and pooling/aggregation operators, then decompress the latent information *Z* into the target *Y* via upsampling and deconvolution operators.

Pix2Pix accomplishes the mapping *G* through generation of *Y* from *X* and *Z*. The original image *X* and target image *Y* are concatenated together, and original image *X* and generated image *G*(*x,z*) are concatenated. Both of these images are passed through the discriminator via the objective:

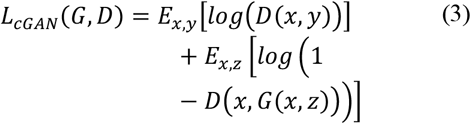

This loss function is supplemented by an L1-Loss that compares the generated image with the target image:

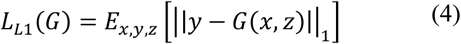

The optimization process is identical to the GAN training process and is performed on a weighted combination of the conditional GAN (cGAN) and L1 objectives.

The CycleGAN model trains two generators, *G*: *X* → *Y* and *F*: *Y* → *X*, and utilizes discriminators, *D_X_* and *D_Y_* for the source and target domains respectively. The CycleGAN objective utilizes a cycle-consistent loss, which assumes that original data *X*, mapped to *Y* via *G*, then mapped back via *F*, should resemble the original data source. This is expressed as:

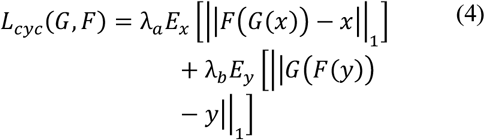

There two weighting terms control the importance of generating from one domain versus the other. The final objective adds the cycle-consistent loss to adversarial losses for the source and target domains:

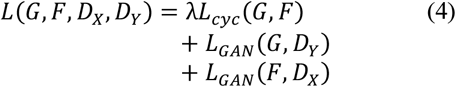

A similar minmax optimization of the objective is used to obtain the ideal parameterization of *G* and *F* to realize the translation from *X* to *Y* when images are unpaired.

### 2 Code and Data Availability

The CycleGAN and Pix2Pix models that were utilized in this study were trained using PyTorch 1.3.0, code was adopted from GitHub repository https://github.com/junyanz/pytorch-CycleGAN-and-pix2pix. Additionally, we provide helper scripts and small test datasets, which will undergo continuous updating, in our GitHub repository: https://github.com/jlevy44/PreliminaryGenerativeHistoPath. PathFlowAI, which was utilized to prepare the data for the H&E to Trichrome analysis is available at: https://github.com/jlevy44/PathFlowAI.

